# Maternal and/or direct supplementation with a combination of a casein hydrolysate and yeast β-glucan on post-weaning performance and intestinal health in the pig

**DOI:** 10.1101/2022.02.23.481664

**Authors:** Eadaoin Conway, John V. O’Doherty, Anindya Mukhopadhya, Alison Dowley, Stafford Vigors, Shane Maher, Marion T. Ryan, Torres Sweeney

**Affiliations:** School of Agriculture and Food Science, University College Dublin, Belfield, Dublin 4, Ireland; School of Veterinary Medicine, University College Dublin, Belfield, Dublin 4, Ireland

**Keywords:** casein hydrolysate, yeast β-glucan, maternal, pig, gut microbiota, intestinal morphology

## Abstract

A 2 × 2 factorial experiment was conducted to investigate the effect of maternal supplementation from day 83 of gestation and/or direct supplementation from weaning of a bovine casein hydrolysate plus a yeast β-glucan (CH-YBG) on pig performance and intestinal health on day ten post-weaning. Twenty cross bred gilts (Large White × Landrace) were randomly assigned to one of two dietary groups (*n* = 10 gilts/group): basal diet (basal sows) and basal diet supplemented with CH-YBG (supplemented sows) from day 83 of gestation until weaning (2g/sow/day). At weaning, 120 pigs (6 pigs/sow) were selected. The two dam groups were further divided, resulting in four experimental groups (10 replicates/group; 3 pigs/pen) as follows: 1) BB (basal sows + basal pigs); 2) BS (basal sows + supplemented pigs); 3) SB (supplemented sows + basal pigs); 4) SS (supplemented sows + supplemented pigs). Supplemented pigs were offered 0.5g CH-YBG/kg of feed for 10 days post-weaning. On day 10 post-weaning, 1 pig/pen was humanely sacrificed and samples were taken from the gastrointestinal tract for analysis. Pigs weaned from supplemented sows (SS, SB) had reduced faecal scores and incidence of diarrhoea (P<0.05) compared to pigs weaned from basal sows (BB, BS), with SS pigs not displaying the transient rise in faecal scores seen in the other three groups from day 3 to day 10 post-weaning (P<0.05). Pigs weaned from supplemented sows had reduced feed intake (P<0.05), improved feed efficiency (P<0.05), increased butyrate concentrations (P<0.05), increased abundance of *Lactobacillus* (P<0.05) and decreased abundance of *Enterobacteriaceae* and *Campylobacteraceae* (P<0.05) compared to pigs weaned from basal sows. In conclusion, a combination of both maternal and direct supplementation with CH-YBG is the desirable feeding strategy to ameliorate the post-weaning challenge in pig production systems.

## 1. Introduction

Traditionally, antibiotic growth promotors and pharmacological levels of zinc oxide were supplemented to pig weaner diets to enhance growth and overcome post-weaning intestinal dysfunction. However due to antimicrobial resistance and environmental contamination, the use of pharmacological levels of zinc oxide is being phased out, with its complete ban imminent in the EU by 2022 (Commission Implementing Decision of 26 June 2017, C (2017) 4529 Final). Therefore, identifying natural feed supplements and management strategies which can alleviate the post-weaning growth check and intestinal dysfunction is of critical importance.

Bovine milk is a rich source of bioactive peptides. However, many of these peptides are biologically inactive in their intact protein and need to be hydrolysed [1]. A moderately hydrolysed (11-16%) bovine casein sample, compared to its parent protein, displayed potent anti-inflammatory activity in both *in vitro* and *ex vivo* systems [2] and also increased *Lactobacillus, Bifidobacterium* and butyrate production while decreasing *Salmonella Typhimurium in vitro* [3]. Supplementation of a casein hydrolysate 5kDa retentate (5kDaR) in an experimental weaned pig model suppressed pro-inflammatory cytokines in the duodenum but not in the jejunum or ileum, suggesting a breakdown of the bioactive molecule during its transit through the small intestine [4]. Natural encapsulating and delivery agents, such as β-glucans, can protect bioactive compounds from digestion in the stomach [5, 6]. In addition, yeast β-glucans themselves exhibit a variety of innate bioactive properties, including anti-inflammatory and immunomodulating properties [7–9]. The potential of combining a combination of a casein 5kDa retentate with a yeast β-glucan as a supplement to the diet of post-weaned pigs was identified, with faecal scores comparable to pigs receiving ZnO supplementation [4].

Traditional measures to ameliorate or reduce post-weaning disorders involved direct supplementation of bioactives to the pig following separation from the dam at the time of weaning. However, we proposed that maternal supplementation with immune supporting natural bioactives during late gestation and lactation would also support the developmental maturation of the digestive tract and thus an alternative approach to preventing the acute challenge of abrupt weaning [10, 11]. Maternal nutrition, particularly during late gestation, plays a critical role in foetal growth and development of neonatal piglets. Maternal supplementation, during the last trimester of gestation, with seaweed bioactives [12] led to improved gut architecture at weaning while also modulating the sow’s faecal microbiota by reducing *Enterbacteriaceae*, leading to the colonisation of beneficial bacteria while reducing *E. coli* in the gastrointestinal tract (GIT) of neonatal pigs [12–14].

Hence, the objectives of this study were: 1) to determine if dietary supplementation with a bovine casein hydrolysate plus a yeast β-glucan (CH-YBG) benefits pig health and performance; and 2) to determine which supplementation period (maternal supplementation from day 83 of gestation to weaning and/or direct supplementation for 10 day’s post-weaning) results in most improved gastrointestinal health and performance. It was hypothesised that pigs exposed to CH-YBG supplement during gestation, lactation and post-weaning would have enhanced performance and improved aspects of intestinal health and immune status, making them more resilient to post-weaning challenges.

## 2. Materials and Methods

All experimental procedures described in the present study were approved under University College Dublin Animal Research Ethics Committee (AREC-17-38-Sweeney) and were conducted in accordance with Irish legislation (SI no. 534/2012) and the EU directive 2010/63/EU for animal experimentation. All efforts were taken to minimise pain and discomfort to the animal while conducting these experiments.

### 2.1 Experimental design and animal management

#### 2.1.1 Experimental diet

The casein hydrolysate (CH) used in this study was produced from the hydrolysis of sodium caseinate (NaCas, ≈ 90% w/w protein, Kerry Food Ingredients, Listowel, Ireland) derived from bovine milk and has previously been reported by Mukhopadhya, Noronha (15). The yeast β-glucan was derived from *Saccharomyces cerevisiae* (Biothera Pharmaceuticals, Inc., Eagan, MI, USA) and the optimum concentrations of a casein hydrolysate and a yeast β-glucan (CH-YBG) were established from previous studies [7, 16].

#### 2.1.2 Gestation and lactation period

A total of 20 pregnant gilts (Large White × Landrace genetic lines) were randomly assigned to one of two dietary groups (*n* = 10 gilts/group): D1) basal gestation/lactation diet (basal sows) and D2) basal gestation/lactation diet plus CH-YBG (supplemented sows) from day 83 of gestation until weaning (day 28). The CH-YBG supplement (2.0 g/sow/d) contained 1.0g CH and 1.0g yeast β-glucan. The ingredient composition of the lactation and gestation diets are presented in Table 1. The gestation diet contained 140 g/kg crude protein, 13.5 MJ/kg of digestible energy and 4.4 g/kg of standardised ileal digestible lysine. The lactation diet contained 190 g/kg of crude protein, 14.5 MJ/kg of digestible energy and 8.5 g/kg of standardised ileal digestible lysine. The amino acid requirements were met relative to lysine [17].

**Table 1.**
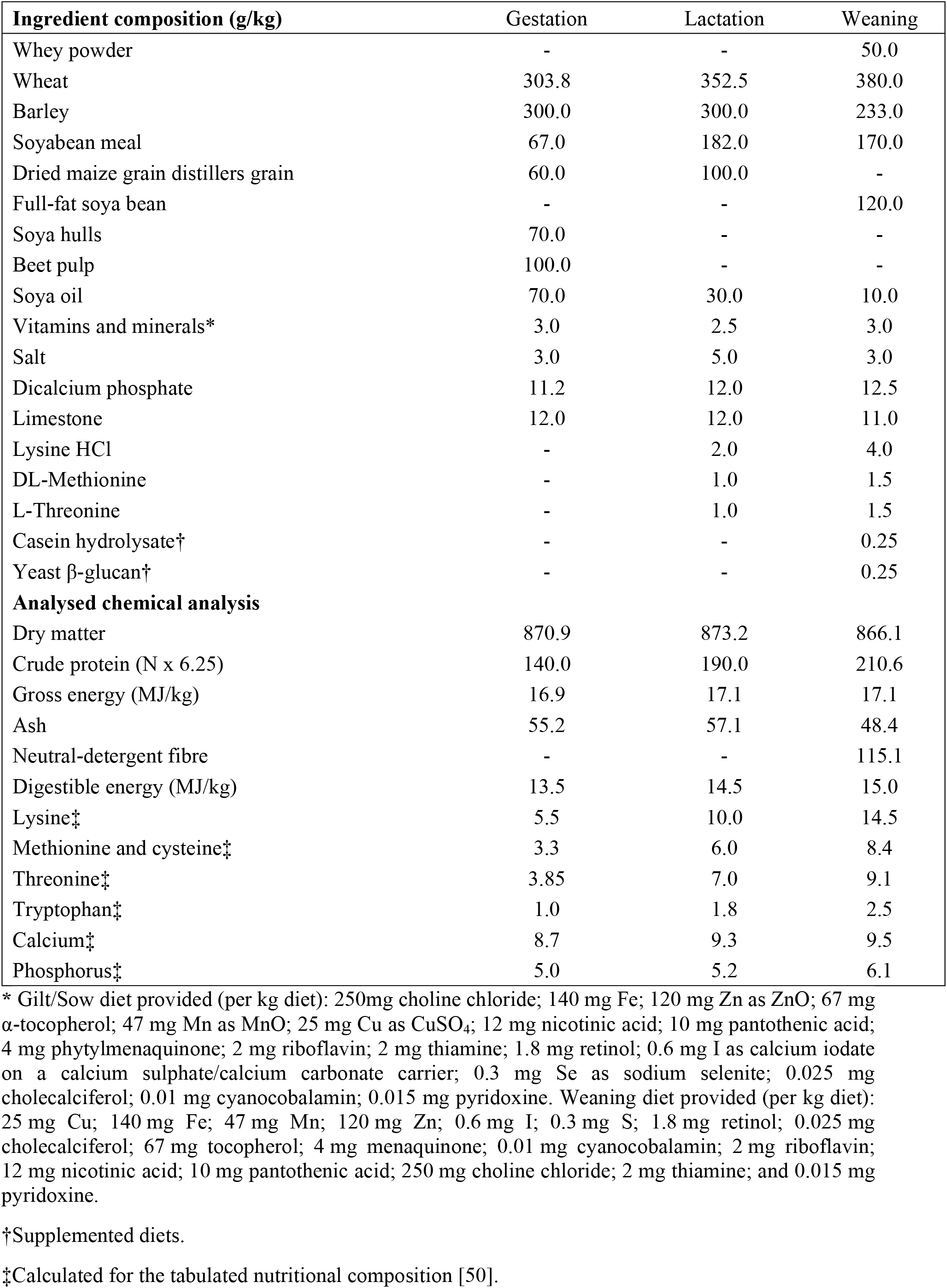
Ingredient and chemical composition of gestation and lactation diet.

| Ingredient composition (g/kg) | Gestation | Lactation | Weaning |
| --- | --- | --- | --- |
| Whey powder | - | - | 50.0 |
| Wheat | 303.8 | 352.5 | 380.0 |
| Barley | 300.0 | 300.0 | 233.0 |
| Soyabean meal | 67.0 | 182.0 | 170.0 |
| Dried maize grain distillers grain | 60.0 | 100.0 | - |
| Full-fat soya bean | - | - | 120.0 |
| Soya hulls | 70.0 | - | - |
| Beet pulp | 100.0 | - | - |
| Soya oil | 70.0 | 30.0 | 10.0 |
| Vitamins and minerals* | 3.0 | 2.5 | 3.0 |
| Salt | 3.0 | 5.0 | 3.0 |
| Dicalcium phosphate | 11.2 | 12.0 | 12.5 |
| Limestone | 12.0 | 12.0 | 11.0 |
| Lysine HCl | - | 2.0 | 4.0 |
| DL-Methionine | - | 1.0 | 1.5 |
| L-Threonine | - | 1.0 | 1.5 |
| Casein hydrolysate† | - | - | 0.25 |
| Yeast β-glucan† | - | - | 0.25 |
| **Analysed chemical analysis** |  |  |  |
| Dry matter | 870.9 | 873.2 | 866.1 |
| Crude protein (N x 6.25) | 140.0 | 190.0 | 210.6 |
| Gross energy (MJ/kg) | 16.9 | 17.1 | 17.1 |
| Ash | 55.2 | 57.1 | 48.4 |
| Neutral-detergent fibre | - | - | 115.1 |
| Digestible energy (MJ/kg) | 13.5 | 14.5 | 15.0 |
| Lysine‡ | 5.5 | 10.0 | 14.5 |
| Methionine and cysteine‡ | 3.3 | 6.0 | 8.4 |
| Threonine‡ | 3.85 | 7.0 | 9.1 |
| Tryptophan‡ | 1.0 | 1.8 | 2.5 |
| Calcium‡ | 8.7 | 9.3 | 9.5 |
| Phosphorus‡ | 5.0 | 5.2 | 6.1 |

\* Gilt/Sow diet provided (per kg diet): 250mg choline chloride; 140 mg Fe; 120 mg Zn as ZnO; 67 mg $\alpha$ -tocopherol; 47 mg Mn as MnO; 25 mg Cu as CuSO<sub>4</sub>; 12 mg nicotinic acid; 10 mg pantothenic acid; 4 mg phytylmenaquinone; 2 mg riboflavin; 2 mg thiamine; 1.8 mg retinol; 0.6 mg I as calcium iodate on a calcium sulphate/calcium carbonate carrier; 0.3 mg Se as sodium selenite; 0.025 mg cholecalciferol; 0.01 mg cyanocobalamin; 0.015 mg pyridoxine. Weaning diet provided (per kg diet): 25 mg Cu; 140 mg Fe; 47 mg Mn; 120 mg Zn; 0.6 mg I; 0.3 mg S; 1.8 mg retinol; 0.025 mg cholecalciferol; 67 mg tocopherol; 4 mg menaquinone; 0.01 mg cyanocobalamin; 2 mg riboflavin; 12 mg nicotinic acid; 10 mg pantothenic acid; 250 mg choline chloride; 2 mg thiamine; and 0.015 mg pyridoxine.
†Supplemented diets.
‡Calculated for the tabulated nutritional composition [50].

From day 83 of gestation to day 110, the gilts were housed in groups of 10. On day 110, gilts were moved to individual farrowing pens (2.2m × 2.4m) with crates, slatted floors and heat pads for the piglets. The gestation house and farrowing room temperature was maintained at 20°C throughout the experiment. The experimental supplement (CH-YBG) was top-dressed on the gestation diet and added to the trough prior to feeding the lactation diet each morning (9am) to ensure consumption. The dams received specific amounts of feed in the following quantities: 2.5 kg/d of gestation diet from day 83 until day 110 of gestation. They were fed 2.0 kg/d of lactation diet from day 111 of gestation until the day of farrowing and then the feed supply increased by 1.0 kg/d until day 3 post-farrowing and by 0.5 kg/d until day 6 post-farrowing. Afterwards, the sows were allowed semi-ad libitum of the diet, which was adjusted for each sow depending on daily intake. The sows were fed in 2 equal meals provided at 9am and 3pm. The sows had ad libitum access to drinking water throughout the experimental period.

#### 2.1.3 Farrowing and piglet management

On the expected farrowing date, fresh sow faecal samples (approximately 10 (SD1.0) g) were collected from each farrowing pen into sterile containers and stored at −20°C for microbial analysis. Farrowing was not induced and were supervised. At parturition, each piglet was individually weighed, and the number of live born piglets were recorded. Six piglets of average birth weight were selected from each litter and ear-tagged shortly after birth. Litter size was adjusted shortly after birth by cross-fostering piglets within dietary groups to ensure that sows nursed a similar number of piglets (12 per sow), and this was maintained throughout the suckling period. No creep feed was offered to the piglets throughout the lactation period, and piglets had no access to the sow’s feed. Piglets received an intramuscular injection of Fedextran (Ferdex 100; Medion Farma Jaja, Indonesia) on day 7 after birth.

#### 2.1.4 Post-weaning period

At weaning, a total of 120 mixed sex pigs (6 pigs/sow) were selected based on their average birth weight. All pigs remained in the same group defined by their dams and subdivided into two groups of 3 pigs, resulting in four experimental groups. The two factors, lactation diet and post-weaning diet, were arranged in a 2 × 2 factorial to provide the four experimental groups that were randomly assigned to replicate pens (3 pigs/pen) as follows: T1) - BB (basal sows + basal pigs); T2) - BS (basal sows + supplemented pigs); T3) - SB (supplemented sows + basal pigs); T4) - SS (supplemented sows + supplemented pigs). Pigs from the BS and SS groups were offered the CH-YBG supplement post-weaning at a rate of 0.5 g/kg of feed.

### 2.2 Starter pig performance

The starter pig performance study measured performance between day 0 and 10 post-weaning, where pigs were supplemented with or without the supplement depending on the experimental group, as described previously. The pigs were housed in groups of 3 (from original sow litter) on fully slatted pens (1.7m × 1.2m). The post-weaning diet contained 210.6 g/kg crude protein, 15.0 MJ/kg digestible energy and 12.5 g/kg standardised ileal digestible lysine. All amino acid requirements were met relative to lysine [17]. All diets were milled on site and fed in meal form for 10 days’ post-weaning. No medication, zinc oxide or other growth-promoting agents were included in the starter diet. The ingredient composition of the post-weaning diet is presented in Table 1. Feed and water were available ad libitum throughout the experimental period. The ambient environmental temperature within the houses was thermostatically controlled. The temperature was maintained at 30°C for the first week and was reduced by 2°C the following week. The pigs were weighed individually on the day of weaning and day 10 post-weaning. Feed intake was recorded per pen to calculate average daily feed intake.

### 2.3 Faecal scoring

From weaning to day 10, faecal scores were assessed twice daily for each individual pen to indicate the presence and severity of diarrhoea. The following scoring system was used to assign faecal scores: 1 = hard, 2 = slightly soft, 3 = soft, partially formed, 4 = loose, semi-liquid, 5 = watery, mucous like [18].

### 2.4 Animal sacrifice and sample collection

On day 10 post-weaning, one pig per pen (selected at birth) was sacrificed following a lethal injection of pentobarbitone sodium (Euthatal Soluion, 200mg/ml; Merial Animal Health, Essex, UK) at a rate of 0.71ml/kg body weight to the cranial vena cava to humanely euthanise the animals. Euthanasia was completed by a competent person in a separate room away from sight and sound of the other pigs. Following this, the entire digestive tract was surgically removed. Sections from the duodenum (10cm from the stomach), the jejunum (60cm from the stomach) and the ileum (15cm from the caecum) were excised and fixed in 10% neutral-buffered formalin. tissue samples were taken from the duodenum, jejunum, ileum and colon to measure the gene expression of cytokines, digestive enzymes, nutrient transporters, mucins, tight junction and appetite regulators using QPCR. Tissue sections of 1 cm^2^ from the duodenum, jejunum, ileum and colon were cut out, emptied by dissecting them along the mesentery and rinsed using sterile phosphate buffer saline (PBS) (Oxoid, Hampshire, UK). The tissue sections were stripped of the overlying smooth tissue before storage in 5ml RNAlater® solution (Applied Biosystems, Foster City, CA, USA) overnight at 4°C. The RNAlater® was removed before storing the samples at −80°C. Digesta samples (approximately 10g) from the caecum and colon were aseptically collected into sterile containers (Sarstedt, Wexford, Ireland) and immediately frozen for subsequent 16s rRNA sequencing and VFA analysis.

### 2.5 Gut morphological analysis

Preserved duodenal, jejunal and ileal tissue samples were prepared using standard paraffin-embedding techniques. The samples were sectioned at a thickness of 5 μm and stained with haematoxylin–eosin. Villus height (VH) and crypt depth (CD) were measured in the stained sections (4× objective) using a light microscope fitted with an image analyser (Image-Pro Plus; Media Cybernetics). Measurements of fifteen well orientated and intact villi and crypts were taken for each segment. The VH was measured from the crypt–villus junction to the tip of the villus, and CD was measured from the crypt– villus junction to the base. Results are expressed as mean VH or CD in μm.

### 2.6 Gene Expression Analysis

#### 2.6.1 RNA Extraction and cDNA synthesis

Total RNA was extracted from jejunal, ileal and colonic tissue using TRIreagent (Sigma-Aldrich, St. Louis, MO, USA) according to the manufacturer’s instructions. The crude RNA extract was further purified using the GenElute™ Mammalian Total RNA Miniprep kit (Sigma-Aldrich) according to the manufacturer’s instructions. A DNase removal step was included using an on-Column DNase 1 Digestion Set (Sigma-Aldrich). The total RNA was quantified using a Nanodrop-ND1000 Spectrophotometer (Thermo Scientific) and the purity was assessed by determining the ratio of the absorbance at 260 nm and 280 nm. The RNA integrity was assessed using an Agilent 2100 Bioanalyzer using an RNA 6000 Nano LabChip kit (Agilent Technologies, Santa Clara, CA, USA). All samples had a 260:280 ratio > 2.0 and an RNA integrity number (RIN) > 8.0. The total RNA (2μg) was reverse transcribed using a High-Capacity cDNA Reverse Transcription Kit (Applied Biosystems) and oligo (dT) primers in a final reaction volume of 40 μL, according to the manufacturer’s instructions. The cDNA was then adjusted to a volume of 360 μL with nuclease-free water.

#### 2.6.2 Quantitative Real-Time PCR (QPCR)

The quantitative PCR (QPCR) reaction mix (20 μl) contained GoTaq qPCR Master Mix (10μl) (Promega, Madison, WI), forward and reverse primers (1.2 μl) (5μM), nuclease-free water (3.8μl) and cDNA (5μl). All QPCR reactions were performed in duplicate on the 7500 ABI Prism Sequence detection System (Applied Biosystems, Foster City, CA). The cycling conditions included a denaturation step of 95°C for 10 mins followed by 40 cycles of 95°C for 15 sec and 60°C for 1 min. All primers were designed using the Primer Express Software (Applied Biosystems, Foster City, CA) and synthesised by MWG Biotech UK Ltd (Milton 227 Keynes, UK) and are presented in Table **2**. Dissociation curves were generated to confirm the specificity of the resulting PCR products. The QPCR assay efficiencies were established by plotting the cycling threshold (CT) values derived from 4-fold serial dilutions of cDNA against their arbitrary quantities and only assays exhibiting 90 –110% efficiency and single products were used in this study. Normalised relative quantities were obtained using the qbase PLUS software (Biogazelle, Ghent, Belgium) from stable reference genes; *YWHAZ, B2M, GAPDH* and *PPIA* for the jejunum and ileum and *YWHAZ*, *B2M* and *PPIA* for the colon. These genes were selected as reference genes based on their M value (<1.5) generated by the GeNorm algorithm within GeNorm. The genes analysed in the current study are as follows: *SLC15A1* (previously known as *PEPT1*); *FABP2; SLC5A1* (previously known as *SGLT1*); *SLC2A1* (previously known as *GLUT1*); *SLC2A2* (previously known as *GLUT2*); *SLC2A5* (previously known as *GLUT5*); *CCK; TNF; CXCL8* (previously known as *IL8*); *IL6; IL10; IL17; IFNG; MUC1; MUC2; TGFB1; TLR4; CLDN1; CLDN3; B2M; GAPDH; PPIA; YWZHAZ*.

**Table 2.**
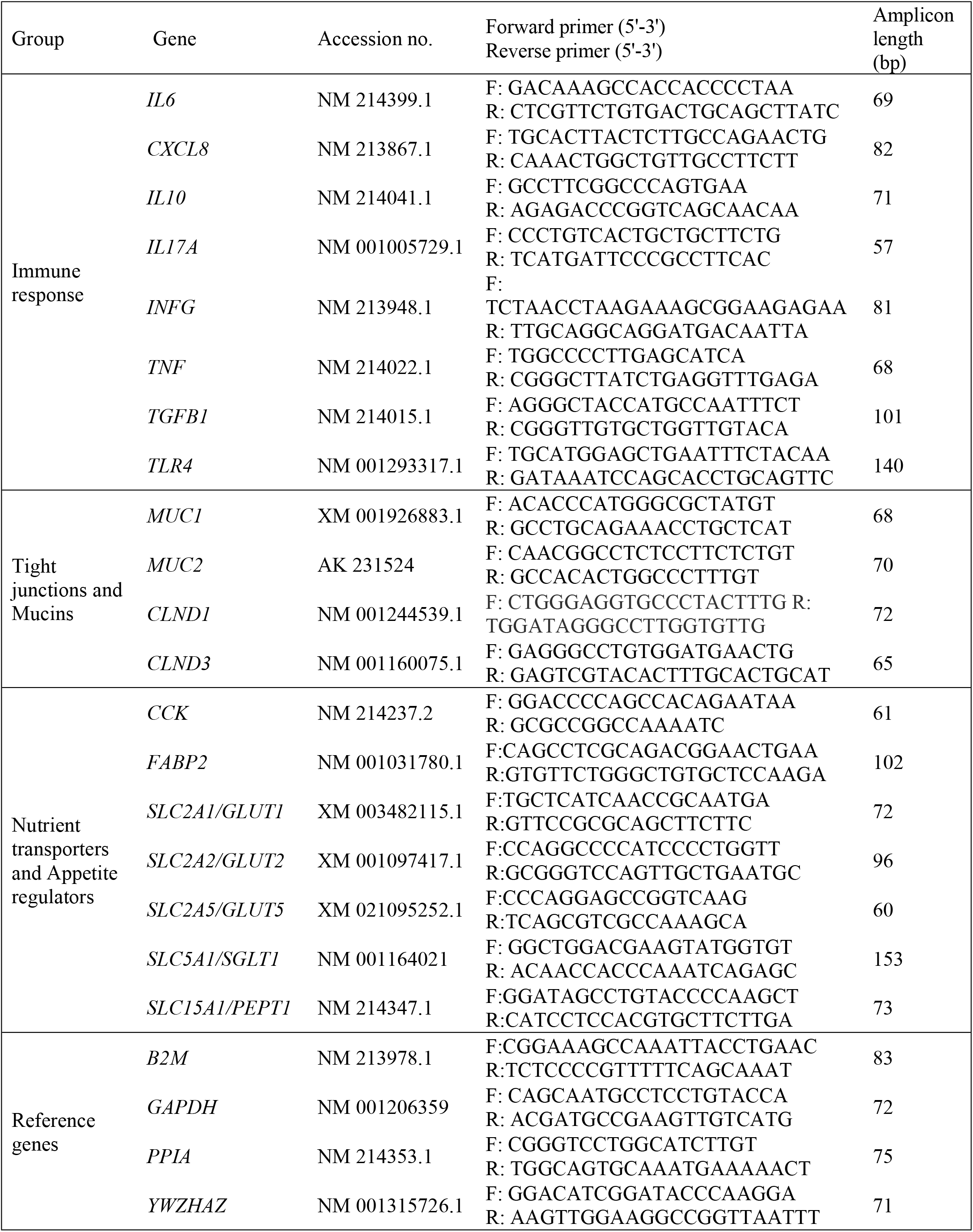
Panel of porcine oligonucleotide primers used for real-time PCR.

### 2.7 Microbial Analysis

#### 2.7.1 Microbial DNA extraction

Microbial genomic DNA was extracted from the pigs colonic digesta samples and the sows faeces using a QIAamp PowerFecal Pro DNA stool kit (Qiagen, West Sussex, UK) according to the manufacturer’s instructions. The quantity and quality of DNA were assessed using a Nanodrop ND-1000 Spectrophotometer (Thermo Scientific, Wilmington, DE).

#### 2.7.2 Illumina sequencing

High-throughput sequencing of the V3-V4 hypervariable region of the bacterial 16S rRNA gene was performed on an Illumina MiSeq platform according to their standard protocols (Eurofins, Genomics, Ebersberg, Germany). Briefly, the V3–V4 region was PCR-amplified using universal primers containing adapter overhang nucleotide sequences for forward and reverse index primers. Amplicons were purified using AMPure XP beads (Beckman Coulter, Indianaopolis, IN) and set up for the index PCR with Nextera XT index primers (Illumina, San Diego, CA). The indexed samples were purified using AMPure XP beads, quantified using a fragment analyser (Agilent, Santa Clara, CA), and equal quantities from each sample were pooled. The resulting pooled library was quantified using the Bioanalyzer 7500 DNA kit (Agilent, Santa Clara, CA) and sequenced using the v3 chemistry (2 × 300 bp paired-end reads).

#### 2.7.3 Bioinformatics and Statistical analysis

The bioinformatic analyses of the resulting sequences was performed by Eurofins Genomics (Eberberg, Germany) using the open source software package (version 1.9.1) Quantitative Insights into Microbial Ecology (Qiime) [19]. All raw reads passing the standard Illumina chastity filter were demultiplexed according to their index sequences (read quality score >30). The primer sequences were clipped from the starts of the raw forward and reverse read. If primer sequences were not perfectly matched, read pairs were removed to retain only high-quality reads. Paired-end reads were then merged if possible, to obtain a single, longer read that covers the full target region using the software FLASH 2.2.00 [20]. Pairs were merged with a minimum overlap size of 10 bp to reduce false-positive merges. The forward read was only retained for the subsequent analysis steps when merging was not possible. Merged reads were quality filtered according to the expected length and known length variations of the V3 – V4 region (ca. 445bp). The ends of retained forward reads were clipped to a total read length of 285 bp to remove low quality bases. Merged and retained reads containing ambiguous bases were discarded. The filter reads (merged and quality clipped retained forward reads) were used for the microbiome profiling. Chimeric reads were identified and removed based on the de-novo algorithm of UCHIME [21] as implemented in the VSEARCH package [22]. The remaining set of high-quality reads was processed using minimum entropy decomposition (**MED**) to partition reads to operational taxonomic units (**OTU**) [23, 24]. DC-MEGABLAST alignments of cluster representative sequences to the NCBI nucleotide sequence database were performed for taxonomic assignment (from phylum to species) of each OTU. A sequence identity of 70% across at least 80% of the representative sequence was the minimal requirement for considering reference sequences. Abundances of bacterial taxonomic units were normalised using lineage-specific copy numbers of the relevant marker genes to improve estimates [25].

The normalised OTU table combined with the phenotype metadata and phylogenetic tree comprised the data matrix. This matrix was then put into the phyloseq package within R (http://www.r-project.org; version 3.5.0). The dynamics of richness and diversity in the pig’s microbiota were computed with the observed, Chaol, ACE, Shannon, Simpson, inverse Simpson and Fisher indices. The Simpson and Shannon indices of diversity account for both richness and evenness parameters. The beta diversity measurements are a measure of separation of the phylogenetic structure of the OTU in one sample compared with all other samples. This was estimated by normalising the data so taxonomic feature counts were comparable across samples. Several distance metrics were considered, in order to calculate the distance matrix of the different multidimensional reduction methods. These included weighted/unweighted UniFrac distance and non-phylogenetic distance metrics (i.e., Bray–Curtis, Jensen–Shannon divergence and Euclidian) using phyloseq in R [26, 27]. Differential abundance testing was performed on tables extracted from the phyloseq object at phylum, family and genus level. The model assessed the effect of ‘group’, with the individual pig being the experimental unit. Ten pigs per group group were used for the statistical analysis of the relative bacterial abundances. The genus *Prevotella-1* belongs to the family *Prevotellaceae* whereas *Prevotella-2* belongs to the family *Paraprevotellaceae*.

### 2.8 Volatile fatty acids analysis

The VFA concentrations in colonic digesta were determined using gas liquid chromatography (GLC) according to the method described by Clarke, Sweeney (28). A 1 g sample of digesta was diluted with distilled water (2·5 × weight of sample) and centrifuged at 1400 × g for 10 min using a Sorvall GLC-2B laboratory centrifuge (DuPont, Wilmington, DE, USA). The supernatant (1 ml) and internal standard (1 ml; 0·05 %o 3-methyl-n-valeric acid in 0·15 M oxalic acid dihydrate) were mixed with 3 ml of distilled water. The mixture was centrifuged at 500 × g for 10 minutes and the supernatant was filtered through 0.45 TFE (polytetrafluoroethylene) syringe filter into a chromatographic sample vial. An injection volume of 1 μl was injected into a Varian 3800 GC (Ontario, Canada) equipped with an EC™ 1000 Grace column (15 m × 0·53 mm I.D) with 1·20 μm film thickness. The temperature programme set was 75–95°C increasing by 3°C/min and 95–200°C increasing by 20°C/min, which was held for 0.50 min. The detector and injector temperature were 280°C and 240°C, respectively, while the total analysis time was 12.42 min.

### 2.9 Feed analysis

The feed samples were milled through a 1 mm screen (Christy and Norris Hammer Mill, Ipswich, UK). The dry matter content of the feed was determined after drying overnight at 104°C. Ash content was determined after ignition of a weighted sample in a muffle furnace (Nabertherm) at 550°C for 6 hr. The gross energy content was determined using an adiabatic bomb calorimeter (Parr Instruments, Illnois, USA). The nitrogen content was determined using the LECO FP 528 instrument (Leco Instruments, UK Ltd). The neutral-detergent fibre content was determined according to Van Soest, Robertson (29) using the Ankom 220 Fibre Analyser (Ankom^tm^ Technology, New York, USA)

### 2.10 Statistical analysis

The data was initially checked for normality using the UNIVARIATE procedure of of SAS® version 9.4 (SAS Institute, Inc.). Growth parameters (ADG, ADFI and gain-to-feed ratio [G:F]), intestinal morphology, gene expression and VFA concentrations and molar proportions were analysed using the PROC GLM procedure of SAS. The model included the two factors – lactation diet and post-weaning diet and their associated interaction. Faecal scores were averaged every two days and analysed using the PROC MIXED procedure of SAS. The model included the two factors – lactation diet and post-weaning diet and their associated two and three way interactions. The incidence of diarrhoea during the first 10 days post-weaning was analysed using PROC Genmod procedure of SAS. The microbiome data was analysed using PROC GLIMMIX. Results are presented as least-square using Benjamini–Hochberg (BH) adjusted P-values. The results are presented as least-square means with their standard errors. When there is no interaction between lactation diet and post-weaning diet, results are presented as main effects. The probability level that denotes significance is P < 0.05.

## 3. Results

### 3.1 Growth performance

The effects of supplementation during gestation/lactation and/or post-weaning on growth performance (ADG, ADFI, weight and G:F) are presented in Table 3. There was no interaction between lactation diet and post-weaning diet on post-weaning performance. Pigs weaned from supplemented sow had reduced (P<0.001) feed intake and increased (P<0.05) gain-to-feed ratio compared to pigs weaned from basal sows. There was no difference (P>0.05) in ADG and day 10 weight across all groups.

**Table 3.** Effect of supplementation of CH-YBG during gestation/lactation and/or post-weaning on pig growth performance and diarrhoea incidence for 10 days post-weaning. (least-square mean values with their standard errors)

|  | Period |  |  |  |  |  | P-value* |  |
| --- | --- | --- | --- | --- | --- | --- | --- | --- |
|  | Lactation |  |  | Post-weaning |  |  |  |  |
|  | Basal | CH-YBG | SEM | Basal | CH-YBG | SEM | LT | PW |
| ADG (g/d) | 166.59 | 153.89 | 7.422 | 161.33 | 159.15 | 7.431 | 0.230 | 0.837 |
| Intake (g/d) | 226.35 | 184.69 | 4.603 | 207.18 | 203.86 | 4.608 | <0.001 | 0.612 |
| Weight (kg) | 8.22 | 8.09 | 0.074 | 8.16 | 8.14 | 0.074 | 0.230 | 0.837 |
| G:F | 0.74 | 0.84 | 0.031 | 0.78 | 0.79 | 0.031 | 0.026 | 0.812 |
| Diarrhoea incidence (D0-10)<br>% † | 22.92 | 6.46 | 0.031 | 21.60 | 6.94 | 0.032 | <0.001 | 0.002 |

CH-YBG, casein hydrolysate plus a yeast β-glucan; ADG, average daily gain; ADFI, average daily feed intake; G:F, gain to feed ratio; LT, lactation diet; PW, post-weaning diet.
\*There was no LT × PW interaction.
†A faecal score of 3 or greater was categorised as diarrhoea.

### 3.2 Faecal scores and diarrhoea incidence

The effects of supplementation during gestation/lactation and/or post-weaning on faecal scores are presented in Figure 1. There was an interaction identified between lactation diet, post-weaning diet and time on faecal scores (P< 0.001). While there was no difference in faecal scores up to day 2 post-weaning (P>0.05), SS pigs did not have the transient rise in faecal scores from day 3 to day 10 post-weaning that was evident in the other three groups (P<0.05).

**Fig 1.**
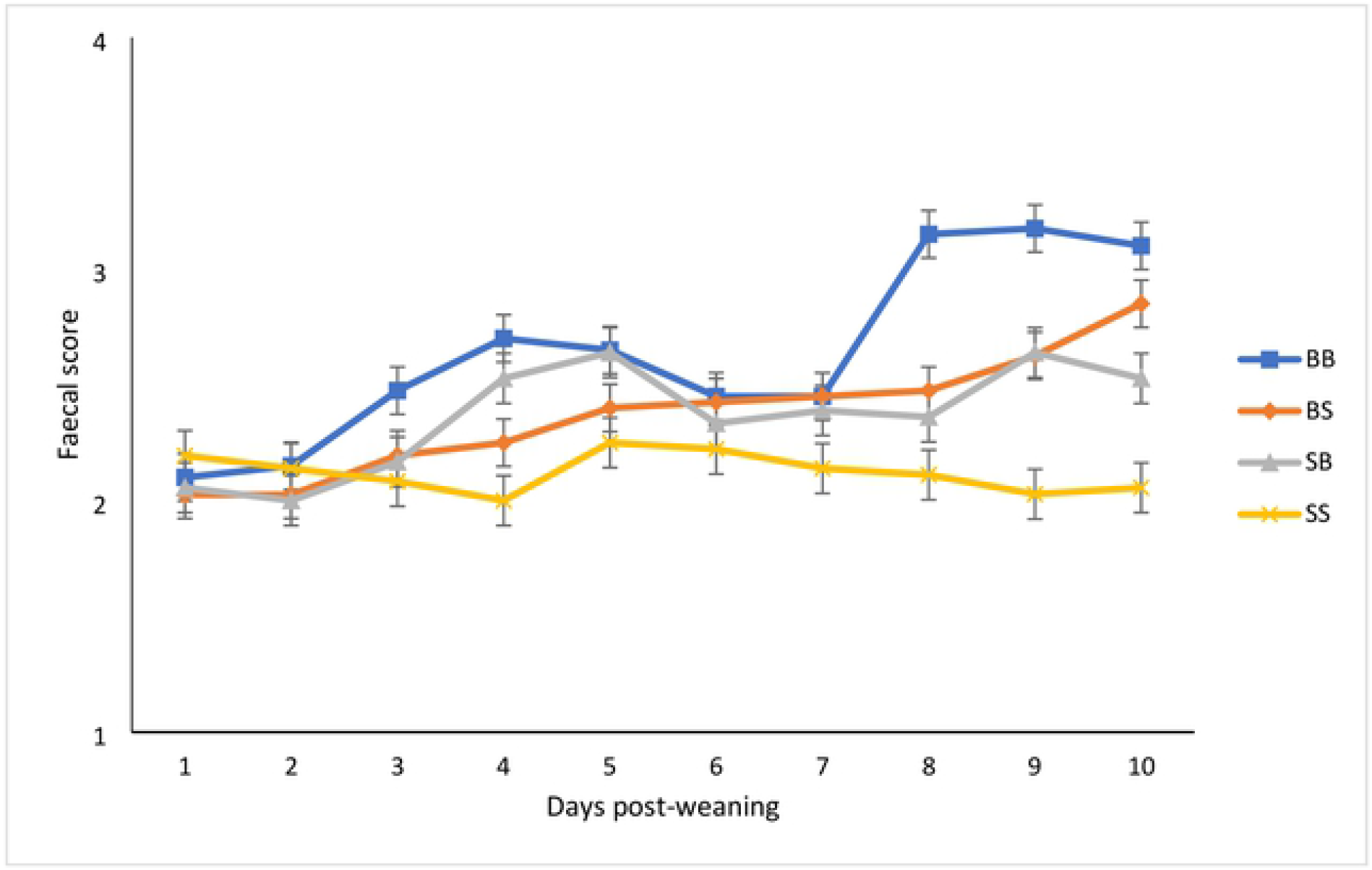
Effect of supplementation of CH-YBG during gestation/lactation and/or post-weaning on faecal scores from day 0-9 post-weaning. BB basal sow, basal pig; BS basal sow, supplemented pig; SB supplemented sow, basal pig; SS supplemented sow, supplemented pig. Values are means, with their standard errors represented by vertical bars. Scale from 1 to 5: 1 = hard, firm faeces; 2 = slightly soft faeces; 3 = soft, partially formed faeces; 4 = loose, semi-liquid faeces; and 5 = watery, mucous-like faeces [18]. Time (p<0.001), lactation diet (p<0.001), post-weaning diet (p<0.001), lactation diet × post-weaning diet (p>0.10), lactation diet × post-weaning diet × time (p<0.001).

The effects of supplementation during gestation/lactation or post-weaning on diarrhoea incidence is presented in Table 3. A faecal score of 3 or greater was categorised as diarrhoea. The diarrhoea incidence for each experimental group was as follows: BB – 34.29%; BS – 14.49%; SB - 12.70% and SS – 3.18%. There was no interaction between lactation diet and post-weaning diet on diarrhoea incidence. Pigs weaned from supplemented sows had a reduced incidence of diarrhoea (P<0.05) compared to pigs weaned from basal sows. Pigs offered the supplement post-weaning had a reduced incidence of diarrhoea (P<0.05) compared to pigs offered a basal diet post-weaning.

### 3.3 Small intestine morphology

The effects of supplementation during gestation/lactation and/or post-weaning on villus height, crypt depth and villus height to crypt depth ratio are presented in Table 4. There was no interaction between lactation diet and post-weaning diet on gut morphology on day 10 post-weaning. There was no difference (P>0.05) in villus height and crypt depth observed in any segments of the small intestine between all experimental groups.

**Table 4.** Effect of supplementation of CH-YBG during gestation/lactation and/or post-weaning on villus height and crypt depth in the small intestine. (least-square mean values with their standard errors)

|  | Period |  |  |  |  |  | P-value* |  |
| --- | --- | --- | --- | --- | --- | --- | --- | --- |
|  | Lactation |  | SEM | Post-weaning |  | SEM |  |  |
|  | Basal | CH-YBG |  | Basal | CH-YBG |  | LT | PW |
| Duodenum |  |  |  |  |  |  |  |  |
| VH (μm) | 299.49 | 306.39 | 14.395 | 293.02 | 312.86 | 14.400 | 0.737 | 0.337 |
| CD (μm) | 129.67 | 126.18 | 5.017 | 124.84 | 131.01 | 5.047 | 0.856 | 0.859 |
| VH:CD | 2.43 | 2.54 | 0.149 | 2.52 | 2.45 | 0.149 | 0.613 | 0.754 |
| Jejunum |  |  |  |  |  |  |  |  |
| VH (μm) | 257.43 | 243.33 | 12.227 | 252.05 | 248.70 | 12.231 | 0.421 | 0.847 |
| CD (μm) | 116.09 | 109.94 | 6.253 | 112.57 | 113.46 | 6.256 | 0.492 | 0.920 |
| VH:CD | 2.24 | 2.37 | 0.182 | 2.37 | 2.24 | 0.182 | 0.632 | 0.600 |
| Ileum |  |  |  |  |  |  |  |  |
| VH (μm) | 265.50 | 265.08 | 12.186 | 256.89 | 273.69 | 12.191 | 0.981 | 0.337 |
| CD (μm) | 114.34 | 119.47 | 4.954 | 115.56 | 118.24 | 4.966 | 0.470 | 0.705 |
| VH:CD | 2.35 | 2.35 | 0.167 | 2.34 | 2.36 | 0.166 | 0.976 | 0.949 |

CH-YBG, casein hydrolysate plus a yeast β-glucan; VH, villus height; CD, crypt depth; LT, lactation diet; PW, post-weaning diet.
\*There was no LT × PW interaction.

### 3.4 Microbiota

#### 3.4.1 Differential Abundance Analysis

##### 3.4.1.1 Phylum

The effects of dietary supplementation on bacterial phyla of the sow on the expected day of farrowing and the pig on day 10 post-weaning are presented in Table 5a and 5b, respectively.

**Table 5a.** Effect of supplementation of CH-YBG during gestation/lactation on differential relative abundance of bacterial taxa at the **phylum** level in sow faeces on expected farrowing date. (least-square mean values with their standard errors)

|  | Treatment* |  | SEM | P-value |
| --- | --- | --- | --- | --- |
|  | Basal | Supplemented |  |  |
| Firmicutes | 82.265 | 93.108 | 2.976 | 0.02 |
| Proteobacteria | 11.344 | 1.687 | 0.724 | <0.001 |
| Actinobacteria | 1.027 | 1.740 | 0.373 | 0.1933 |
| Spirochaetes | 4.147 | 2.851 | 0.588 | 0.1463 |
| Synergistetes | 0.008 | 0.189 | 0.086 | 0.3752 |
| Tenericutes | 0.742 | 0.205 | 0.205 | 0.1321 |

\*A total of ten replicates were used per treatment.

**Table 5b.** Effect of supplementation of CH-YBG during gestation/lactation and/or post-weaning on differential relative abundance of bacterial taxa at the **phylum** level in pig colonic digesta on day 10 post-weaning. (least-square mean values with their standard errors)

|  | Treatment* |  |  |  | SEM | P-value |  |  |
| --- | --- | --- | --- | --- | --- | --- | --- | --- |
|  | BB | BS | SB | SS |  | LT | PW | LT x PW |
| **Colon** |  |  |  |  |  |  |  |  |
| Firmicutes | 66.731 | 68.638 | 82.102 | 83.441 | 3.874 | 0.001 | 0.674 | 0.909 |
| Bacteroidetes | 18.091 | 21.347 | 11.461 | 12.584 | 1.767 | 0.001 | 0.281 | 0.760 |
| Proteobacteria | 11.944 | 5.982 | 3.519 | 1.802 | 1.020 | <0.001 | 0.010 | 0.962 |
| Spirochaetes | 0.805 | 2.345 | 1.531 | 0.505 | 0.489 | 0.351 | 0.967 | 0.032 |

BB basal sow, basal pig; BS basal sow, supplemented pig; SB supplemented sow, basal pig; SS supplemented sow, supplemented pig; LT, lactation diet; PW, post-weaning diet
\*A total of ten replicates were used per treatment.

###### Sow

Supplementation of CH-YBG to the sow increased the abundance of Firmicutes (P<0.05) and decreased the abundance of Proteobacteria (P<0.001) compared to basal sows. There was no difference in the abundance of Actinobacteria, Spirochaetes or Tenericutes between dietary groups.

###### Pig colonic contents

Pigs weaned from supplemented sows had increased abundance of Firmicutes (P<0.05) and decreased abundance of Bacteroidetes and Proteobacteria (P<0.05) compared to pigs weaned from basal sows. Pigs offered the supplement post-weaning had decreased abundance of Proteobacteria (P<0.05) compared to pigs offered a basal diet post-weaning.

There was an interaction identified between lactation diet and post-weaning diet on the abundance of Spirochaetes (P<0.05). BS pigs had increased abundance of Spirochaetes compared to BB pigs. However, there was no effect of supplementation post-weaning when pigs were weaned from supplemented sows.

##### 3.4.1.2 Family

The effects of dietary supplementation on bacterial family of the sow on the expected day of farrowing and the pig on day 10 post-weaning are presented in Table 6a and 6b, respectively.

**Table 6a.** Effect of supplementation of CH-YBG during gestation/lactation and/or post-weaning on differential relative abundance of bacterial taxa at the **family** level in sow faeces on expected farrowing date. (least-square mean values with their standard errors)

|  | Treatment* |  | SEM | P-value |
| --- | --- | --- | --- | --- |
|  | Basal | Supplemented |  |  |
| Lachnospiraceae | 14.057 | 18.725 | 1.286 | 0.02 |
| Clostridiaceae | 25.588 | 18.536 | 1.480 | 0.004 |
| Lactobacillaceae | 5.037 | 21.343 | 1.108 | <0.001 |
| Christensenellaceae | 12.312 | 8.930 | 1.027 | 0.04 |
| Planococcaceae | 1.511 | 0.346 | 0.283 | 0.03 |
| Moraxellaceae | 9.072 | 0.552 | 0.578 | <0.001 |

\*A total of ten replicates were used per treatment

**Table 6b.**
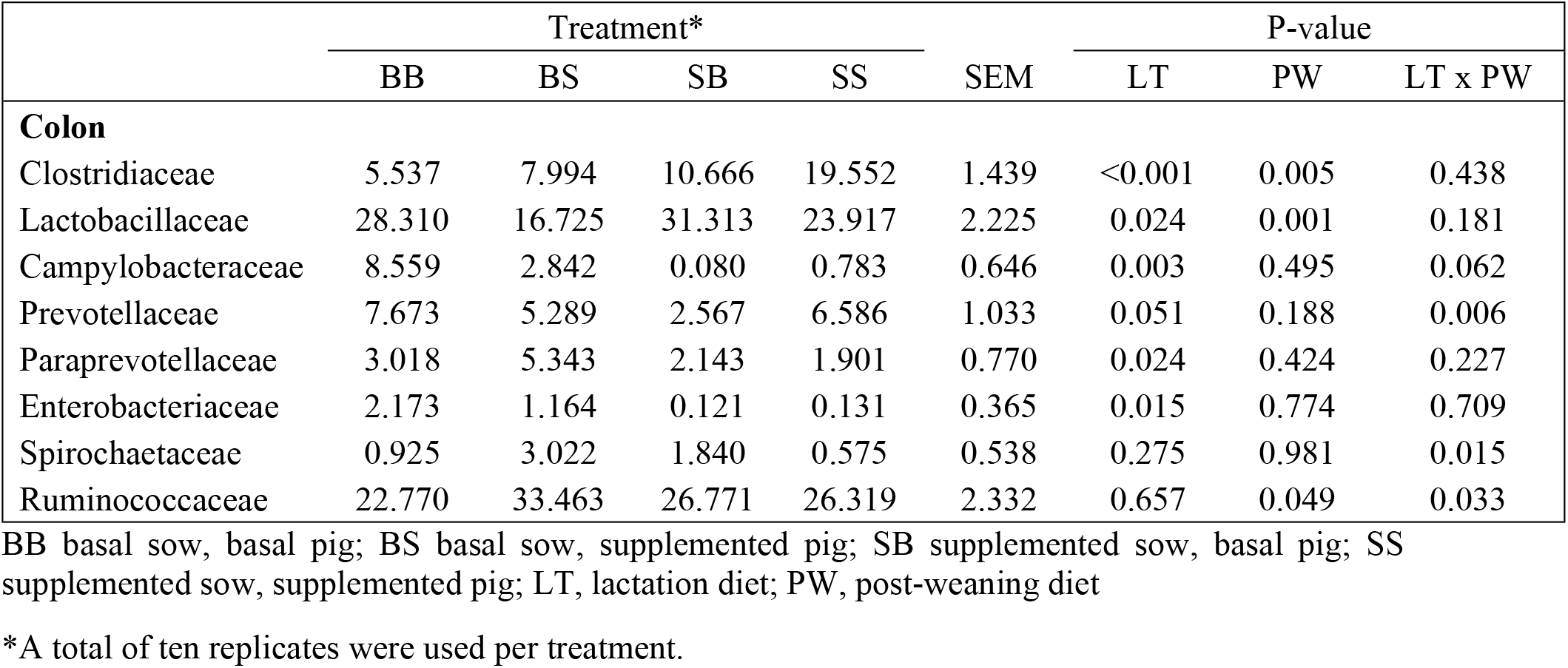
Effect of supplementation of CH-YBG during gestation/lactation and/or post-weaning on differential relative abundance of bacterial taxa at the **family** level in pig colonic digesta on day 10 post-weaning. (least-square mean values with their standard errors)

|  | Treatment* |  |  |  | SEM | P-value |  |  |
| --- | --- | --- | --- | --- | --- | --- | --- | --- |
|  | BB | BS | SB | SS |  | LT | PW | LT x PW |
| **Colon** |  |  |  |  |  |  |  |  |
| Clostridiaceae | 5.537 | 7.994 | 10.666 | 19.552 | 1.439 | <0.001 | 0.005 | 0.438 |
| Lactobacillaceae | 28.310 | 16.725 | 31.313 | 23.917 | 2.225 | 0.024 | 0.001 | 0.181 |
| Campylobacteraceae | 8.559 | 2.842 | 0.080 | 0.783 | 0.646 | 0.003 | 0.495 | 0.062 |
| Prevotellaceae | 7.673 | 5.289 | 2.567 | 6.586 | 1.033 | 0.051 | 0.188 | 0.006 |
| Paraprevotellaceae | 3.018 | 5.343 | 2.143 | 1.901 | 0.770 | 0.024 | 0.424 | 0.227 |
| Enterobacteriaceae | 2.173 | 1.164 | 0.121 | 0.131 | 0.365 | 0.015 | 0.774 | 0.709 |
| Spirochaetaceae | 0.925 | 3.022 | 1.840 | 0.575 | 0.538 | 0.275 | 0.981 | 0.015 |
| Ruminococcaceae | 22.770 | 33.463 | 26.771 | 26.319 | 2.332 | 0.657 | 0.049 | 0.033 |

BB basal sow, basal pig; BS basal sow, supplemented pig; SB supplemented sow, basal pig; SS supplemented sow, supplemented pig; LT, lactation diet; PW, post-weaning diet
\*A total of ten replicates were used per treatment.

###### Sow

Supplementation of CH-YBG to the sow, increased the abundance of *Lactobacillaceae* (P<0.001) and *Lachnospiraceae* (P<0.05) and decreased the abundance of *Clostridiaceae* (P<0.05), *Christensenellaceae* (P<0.05), *Planococcaceae* (P<0.05) and *Moraxellaceae* (P<0.001) compared to basal sows.

###### Pig colonic content

Pigs weaned from supplemented sows had increased abundance of *Lactobacillaceae* and *Clostridiaceae* (P<0.05) and decreased abundance of *Campylobacteraceae, Paraprevotellaceae* and *Enterobacteriaceae* (P<.0.05) compared to pigs weaned from basal sows. Pigs offered the supplement post-weaning had increased abundance of *Clostridiaceae* and decreased *Lactobacillaceae* (P<0.05) compared to pigs offered a basal diet post-weaning.

There was an interaction identified between lactation diet and post-weaning diet on the abundance of *Prevotellaceae* (P<0.05), *Spirochaetaceae* (P<0.05) and *Ruminococcaceae* (P<0.05). SB pigs had decreased abundance of *Prevotellaceae* compared to BB pigs. However, there was no effect of maternal supplementation on *Prevotellaceae* when pigs were offered the supplement post-weaning. BS pigs had increased abundance of *Spirochaetaceae* and *Ruminococcaceae* compared to BB pigs. However, there was no effect of supplementation post-weaning on *Spirochaetaceae* and *Ruminococcaceae* when pigs were weaned from supplemented sows.

##### 3.4.1.3 Genus level

The effects of dietary supplementation on bacterial genus of the sow on the expected day of farrowing and the pig on day 10 post-weaning are presented in Table 7a and 7b, respectively.

###### Sow

Supplementation of CH-YBG to the sow, increased the abundance of *Lactobacillus* (P<0.001), *Gemmiger* (P<0.05), *Niameybacter* (P<0.05) and *Metabacterium* (P<0.001) and decreased the abundance of *Clostridium* (P<0.05), *Kineothrix* (P<0.05), *Christensenella* (P<0.05) and *Acinetobacter* (P<0.001).

**Table 7a.** Effect of supplementation of CH-YBG during gestation/lactation and/or post-weaning on differential relative abundance of bacterial taxa at the **genus** level in sow faeces on expected farrowing date. (least-square mean values with their standard errors)

|  | Treatment* |  | SEM | P-value |
| --- | --- | --- | --- | --- |
|  | Basal | Supplemented |  |  |
| Lactobacillus | 4.975 | 21.083 | 1.102 | <0.001 |
| Clostridium | 25.562 | 17.898 | 1.467 | 0.002 |
| Gemmiger | 0.056 | 1.691 | 0.252 | 0.017 |
| Kineothrix | 3.526 | 1.424 | 0.482 | 0.012 |
| Christensenella | 11.128 | 7.497 | 0.959 | 0.018 |
| Niameybacter | 0.111 | 1.765 | 0.272 | 0.009 |
| Metabacterium | 0.298 | 4.201 | 0.424 | <0.001 |
| Acinetobacter | 9.431 | 0.547 | 0.586 | <0.001 |

\*A total of ten replicates were used per treatment

###### Pig colonic content

Pigs weaned from supplemented sows had increased abundance of *Lactobacillus* (P<0.05) and decreased abundance of *Prevotella-1, Prevotella-2* and *Ruminococcus* (P<0.05) compared to pigs weaned from basal sows. Pigs offered the supplement post-weaning had increased abundance of *Prevotella-2* (P<0.05) and decreased abundance of *Lactobacillus* (P<0.05) compared to pigs offered a basal diet post-weaning.

There was an interaction identified between lactation diet and post-weaning diet on the abundance of *Campylobacter* (P<0.05), *Treponema* (P<0.05) and *Lachnospira* (P<0.05). BS pigs had decreased abundance of *Campylobacter* and increased abundance of *Treponema* compared to BB pigs. However, there was no effect of supplementation post-weaning when pigs were weaned from supplemented sows. SB pigs had increased abundance of *Lachnospira* compared to BB pigs. However, there was no effect of maternal supplementation when pigs were offered the supplement post-weaning.

**Table 7b.** Effect of supplementation of CH-YBG during gestation/lactation and/or post-weaning on differential relative abundance of bacterial taxa at the **genus** level in pig colonic digesta on day 10 post-weaning. (least-square mean values with their standard errors)

|  | Treatment* |  |  |  | SEM | P-value |  |  |
| --- | --- | --- | --- | --- | --- | --- | --- | --- |
|  | BB | BS | SB | SS |  | LT | PW | LT x PW |
| Lactobacillus | 45.526 | 33.565 | 55.967 | 51.153 | 3.038 | <0.001 | 0.009 | 0.127 |
| Campylobacter | 12.409 | 5.383 | 0.152 | 1.532 | 0.835 | <0.001 | 0.246 | 0.021 |
| Prevotella-1 | 12.372 | 10.641 | 5.921 | 8.634 | 1.397 | 0.009 | 0.480 | 0.112 |
| Prevotella-2 | 3.649 | 8.394 | 2.541 | 3.299 | 0.919 | 0.014 | 0.032 | 0.238 |
| Treponema | 1.507 | 5.829 | 3.653 | 1.236 | 0.745 | 0.299 | 0.669 | 0.001 |
| Ruminococcus | 6.015 | 8.283 | 5.494 | 3.799 | 1.076 | 0.037 | 0.900 | 0.091 |
| Lachnospira | 0.682 | 2.398 | 3.277 | 2.083 | 0.629 | 0.068 | 0.287 | 0.032 |

BB basal sow, basal pig; BS basal sow, supplemented pig; SB supplemented sow, basal pig; SS supplemented sow, supplemented pig; LT, lactation diet; PW, post-weaning diet
\*A total of ten replicates were used per treatment.

### 3.5 VFAs

The effects of supplementation during gestation/lactation and/or post-weaning on the concentrations and molar proportions of colonic VFAs are presented in Table 8. There was no interaction between lactation diet and post-weaning diet on VFA production. Pigs weaned from supplemented sows had increased (P<0.05) concentrations of butyrate, isobutyrate and valerate compared to pigs weaned from basal sows. The molar proportions of butyrate and valerate was increased (P<0.05) and acetate decreased (P<0.05) in pigs from supplemented sows compared to pigs weaned from basal sows.

**Table 8.** Effect of supplementation of CH-YBG during gestation/lactation and/or post-weaning on total and molar proportions of volatile fatty acids (VFA) in colonic digesta. (least-square mean values with their standard errors)

|  | Period |  |  |  |  |  | P-value* |  |
| --- | --- | --- | --- | --- | --- | --- | --- | --- |
|  | Lactation |  | SEM | Post-weaning |  |  |  |  |
|  | Basal | CH-YBG |  | Basal | CH-YBG | SEM | LT | PW |
| VFA (mmol/l digesta) |  |  |  |  |  |  |  |  |
| Total | 141.00 | 144.54 | 5.783 | 140.67 | 144.87 | 5.785 | 0.668 | 0.612 |
| Acetate | 109.10 | 107.80 | 4.404 | 108.10 | 108.80 | 4.406 | 0.835 | 0.910 |
| Propionate | 16.46 | 17.34 | 0.897 | 16.46 | 17.34 | 0.898 | 0.492 | 0.493 |
| Butyrate | 9.83 | 12.92 | 0.959 | 10.23 | 12.52 | 0.960 | 0.029 | 0.102 |
| Isobutyrate | 0.93 | 1.36 | 0.126 | 1.11 | 1.18 | 0.126 | 0.020 | 0.678 |
| Isovalerate | 3.14 | 3.64 | 0.292 | 3.33 | 3.45 | 0.292 | 0.232 | 0.781 |
| Valerate | 1.54 | 2.06 | 0.121 | 1.66 | 1.93 | 0.121 | 0.005 | 0.124 |
| Molar proportions |  |  |  |  |  |  |  |  |
| Acetate | 0.77 | 0.75 | 0.007 | 0.77 | 0.75 | 0.007 | 0.008 | 0.123 |
| Propionate | 0.12 | 0.12 | 0.004 | 0.12 | 0.12 | 0.004 | 0.699 | 0.706 |
| Butyrate | 0.07 | 0.09 | 0.004 | 0.07 | 0.08 | 0.004 | 0.001 | 0.067 |
| Isobutyrate | 0.01 | 0.01 | 0.001 | 0.01 | 0.01 | 0.001 | 0.078 | 0.748 |
| Isovalerate | 0.02 | 0.03 | 0.003 | 0.02 | 0.03 | 0.003 | 0.496 | 0.765 |
| Valerate | 0.01 | 0.01 | 0.001 | 0.01 | 0.01 | 0.001 | 0.003 | 0.156 |

CH-YBG, casein hydrolysate plus a yeast β-glucan; VFA, volatile fatty acids; LT, lactation diet; PW, post-weaning diet.
\*There was no LT × PW interaction.

### 3.6 Gene expression

The effects of supplementation during gestation/lactation and/or post-weaning on the expression of genes related to appetite regulation, nutrient digestion and absorption, immunity and mucosal barrier function in the jejunum, ileum and colon are presented in Table 9a, Table 9b and Table 9c, respectively. There was no interaction between lactation diet and post-weaning diet on gene expression.

**Table 9a.** Effect of supplementation of CH-YBG during gestation/lactation and/or post-weaning on the expression appetite regulators, tight junctions, nutrient transporters, cytokines and mucins in the jejunum. (least-square mean values with their standard errors)

|  | Period |  |  |  |  |  | P-value* |  |
| --- | --- | --- | --- | --- | --- | --- | --- | --- |
|  | Lactation |  | SEM | Post-weaning |  | SEM |  |  |
|  | Basal | CH-YBG |  | Basal | CH-YBG |  | LT | PW |
| **Appetite regulators** |  |  |  |  |  |  |  |  |
| *CCK* | 1.190 | 1.097 | 0.177 | 1.116 | 1.171 | 0.177 | 0.713 | 0.828 |
| **Tight junctions** |  |  |  |  |  |  |  |  |
| *CLDN1* | 0.739 | 1.403 | 0.163 | 0.990 | 1.151 | 0.163 | 0.007 | 0.490 |
| *CLDN3* | 1.214 | 1.136 | 0.163 | 1.304 | 1.045 | 0.163 | 0.739 | 0.269 |
| **Nutrient transporters** |  |  |  |  |  |  |  |  |
| *SLC2A1/GLUT1* | 0.912 | 1.201 | 0.104 | 1.202 | 0.911 | 0.104 | 0.058 | 0.056 |
| *SLC2A2/GLUT2* | 1.419 | 1.041 | 0.195 | 1.445 | 1.015 | 0.195 | 0.180 | 0.129 |
| *SLC2A5/GLUT5* | 1.426 | 1.019 | 0.196 | 1.279 | 1.166 | 0.196 | 0.153 | 0.687 |
| *SLC15A1/PEPT1* | 1.534 | 0.977 | 0.200 | 1.379 | 1.132 | 0.200 | 0.058 | 0.390 |
| *SLC5A1/SGLT1* | 1.217 | 0.984 | 0.100 | 1.205 | 0.997 | 0.100 | 0.108 | 0.149 |
| *FABP2* | 1.801 | 1.112 | 0.218 | 1.635 | 1.277 | 0.218 | 0.032 | 0.255 |
| **Cytokines** |  |  |  |  |  |  |  |  |
| *TGFB* | 1.069 | 1.316 | 0.175 | 1.127 | 1.258 | 0.175 | 0.326 | 0.602 |
| *IFNG* | 1.027 | 1.473 | 0.213 | 1.410 | 1.090 | 0.214 | 0.149 | 0.298 |
| *TNF* | 1.102 | 1.682 | 0.220 | 1.063 | 1.721 | 0.220 | 0.071 | 0.042 |
| *TLR4* | 0.982 | 1.206 | 0.116 | 1.165 | 1.022 | 0.116 | 0.181 | 0.388 |
| *IL10* | 1.061 | 1.654 | 0.254 | 1.113 | 1.602 | 0.254 | 0.107 | 0.183 |
| *IL6* | 1.294 | 2.071 | 0.336 | 1.407 | 1.958 | 0.336 | 0.112 | 0.255 |
| *IL17* | 1.677 | 0.964 | 0.280 | 1.129 | 1.512 | 0.280 | 0.081 | 0.340 |
| *CXCL8/IL8* | 1.425 | 1.013 | 0.208 | 1.309 | 1.130 | 0.208 | 0.171 | 0.548 |
| **Mucins** |  |  |  |  |  |  |  |  |
| *MUC1* | 1.170 | 0.855 | 0.118 | 1.090 | 0.935 | 0.118 | 0.068 | 0.358 |
| *MUC2* | 1.704 | 1.154 | 0.270 | 1.458 | 1.400 | 0.270 | 0.160 | 0.879 |

*CCK*, cholecystokinin; *CLND1*, claudin 1; *CLDN3*, claudin 3; *SLC2A1/GLUT1*, glucose transporter 1; *SLC2A2/GLUT2*, glucose transporter 2; *SLC2A5/GLUT5*, glucose transporter 5; *SLC15A1/PEPT1*, peptide transporter 1; *SLC5A1/SGLT1*, sodium glucose linked transporter 1; *FABP2*, fatty acid binding protein 2; TGF- $\beta$ , transforming growth factor beta; *IFNG*, interferon gamma; *TNF*, tumor necrosis factor alpha; *TLR4*, toll like receptor 4; *IL10*, interleukin 10; *IL6*, interleukin 6; *IL17*, interleukin 17; *CXCL8*, interleukin 8; *MUC1*, mucin 1; *MUC2*, mucin 2; CH-YBG, casein hydrolysate plus a yeast β-glucan; LT, lactation diet; PW, post-weaning diet.
\*There was no LT × PW interaction.

**Table 9b.**
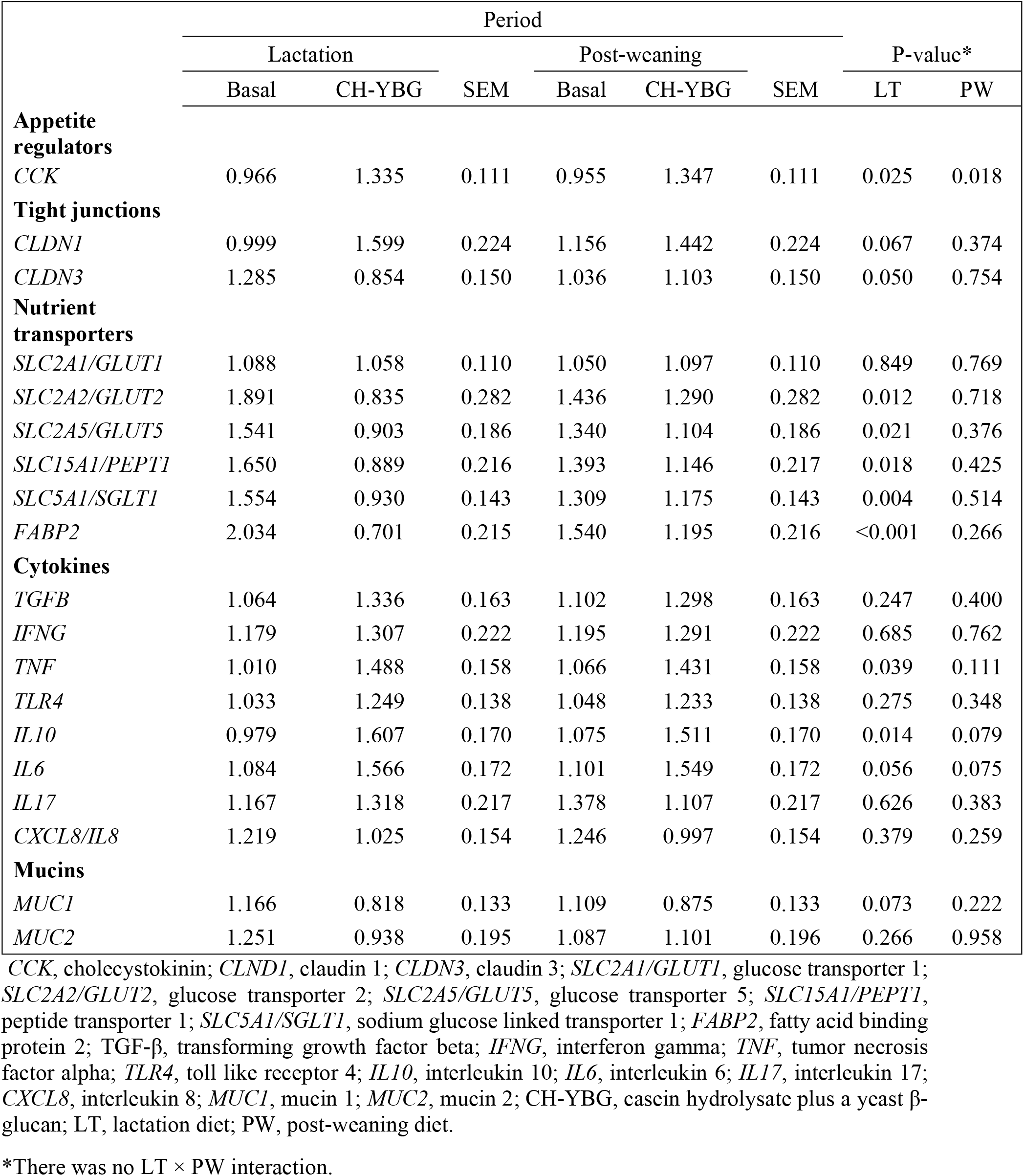
Effect of supplementation of CH-YBG during gestation/lactation and/or post-weaning on the expression appetite regulators, tight junctions, nutrient transporters, cytokines and mucins in the ileum. (least-square mean values with their standard errors)

**Table 9c.** Effect of supplementation of CH-YBG during gestation/lactation and/or post-weaning on the expression appetite regulators, tight junctions, cytokines and mucins in the colon (least-square mean values with their standard errors)

|  | Period |  |  |  |  |  | P-value* |  |
| --- | --- | --- | --- | --- | --- | --- | --- | --- |
|  | Lactation |  |  | Post-weaning |  |  |  |  |
|  | Basal | CH-YBG | SEM | Basal | CH-YBG | SEM | LT | PW |
| **Cytokines** |  |  |  |  |  |  |  |  |
| *TGFB* | 0.951 | 1.113 | 0.084 | 0.975 | 1.090 | 0.084 | 0.183 | 0.341 |
| *IFNG* | 1.125 | 0.993 | 0.164 | 0.989 | 1.130 | 0.164 | 0.575 | 0.547 |
| *TNF* | 0.919 | 1.002 | 0.077 | 0.926 | 0.995 | 0.077 | 0.453 | 0.527 |
| *TLR4* | 0.873 | 1.172 | 0.074 | 0.957 | 1.088 | 0.074 | 0.008 | 0.222 |
| *IL10* | 0.932 | 1.175 | 0.181 | 0.900 | 1.206 | 0.181 | 0.348 | 0.240 |
| *IL6* | 0.923 | 1.291 | 0.103 | 1.140 | 1.074 | 0.103 | 0.016 | 0.649 |
| *IL17* | 1.123 | 1.086 | 0.123 | 1.076 | 1.133 | 0.123 | 0.831 | 0.746 |
| *CXCL8/IL8* | 0.947 | 1.124 | 0.102 | 0.999 | 1.072 | 0.102 | 0.229 | 0.613 |
| **Mucins** |  |  |  |  |  |  |  |  |
| *MUC1* | 0.984 | 1.031 | 0.103 | 1.060 | 0.954 | 0.104 | 0.749 | 0.476 |
| *MUC2* | 1.115 | 1.028 | 0.102 | 1.039 | 1.104 | 0.102 | 0.552 | 0.653 |

TGF- $\beta$ , transforming growth factor beta; *IFNG*, interferon gamma; *TNF*, tumor necrosis factor alpha; *TLR4*, toll like receptor 4; *IL10*, interleukin 10; *IL6*, interleukin 6; *IL17*, interleukin 17; *CXCL8*, interleukin 8; *MUC1*, mucin 1; *MUC2*, mucin 2; CH-YBG, casein hydrolysate plus a yeast β-glucan; LT, lactation diet; PW, post-weaning diet.
\*There was no LT × PW interaction

#### 3.6.1 Nutrient transporter gene expression

##### Jejunum

Pigs weaned from supplemented sows had an upregulation of fatty acid binding protein 2 (*FABP2*, P<0.05) compared with pigs weaned from basal sows.

##### Ileum

Pigs weaned from supplemented sows had a down regulation of glucose transporter 2 (*GLUT2*, P<0.05), glucose transporter 5 (*GLUT5*, P<0.05), peptide transporter 1 (*PEPT1*, P<0.05), sodium glucose linked transporter 1 (*SGLT1*, P<0.05) and *FABP2* (P<0.05) compared to pigs weaned from basal sows.

#### 3.6.2 Appetite regulator gene expression

##### Ileum

Pigs weaned from supplemented sows or pigs offered the supplement post-weaning had an upregulation of cholecystokinin (*CCK*, P<0.05) compared to pigs weaned from basal sows or pigs offered a basal diet post-weaning.

#### 3.6.3 Tight junction and immune marker gene expression

##### Jejunum

Pigs weaned from supplemented sows had an upregulation of claudin 1 (*CLDN1*, P<0.05) compared to pigs weaned from basal sows. Pigs offered the supplement post-weaning had an upregulation of tumor necrosis factor (*TNF*, P<0.05) compared to pigs offered a basal diet post-weaning.

##### Ileum

Pigs weaned from supplemented sows had an upregulation of *TNF* (P<0.05) and interleukin 10 (*IL10*, P<0.05) compared to pigs weaned from basal sows.

##### Colon

Pigs weaned from supplemented sows had an upregulation of toll like receptor 4 (*TLR4*, P<0.05) and interleukin 6 (*IL6*, P<0.05) compared to pigs from basal sows.

## 4. Discussion

It was hypothesised that pigs exposed to the CH-YBG supplement during gestation, lactation and post-weaning would have enhanced performance and improved aspects of intestinal health and immune status, thus making them more resilient to post-weaning challenges. Previous research has suggested that post-weaning supplementation of CH-YBG is a suitable feeding strategy to combat the post-weaning challenge [4]. However, the current study demonstrates that maternal supplementation along with post-weaning supplementation enhanced intestinal health and in fact suppresses the transient rise in faecal scores and diarrhoeal incidence that is typically evident immediately post-weaning. Maternal supplementation also led to increased feed efficiency and VFA production, increased abundance of *Lactobacillus* and a reduction in *Enterobacteriaceae* and *Campylobacter*. Concurrently, genes involved in barrier function were upregulated.

Consistent with previous studies, Firmicutes, Proteobacteria and Bacteroidetes were the three most dominant phyla in the weaned pig [30–32]. Modulation of the pig’s microbiota through maternal supplementation with CH-YBG is evident in this study as both supplemented sows and their offspring exhibited increased Firmicutes and decreased Proteobacteria. Within the phylum Firmicutes, supplemented sows had a greater abundance of the family *Lactobacillaceae* and genus *Lactobacillus* compared to basal sows - an effect that was also observed in their offspring, as pigs weaned from supplemented sows (SB, SS) had increased abundance of *Lactobacillaceae* and *Lactobacillus* compared to pigs weaned from basal sows (BB, BS). Bacteria from the genus *Lactobacillus* have the capacity to lower colonic pH through the production of lactic acid thus inhibiting the growth of pathogenic bacteria [33]. Maternal supplementation led to a reduction in the abundance of *Enterobacteriacea* while maternal or post-weaning supplementation reduced the abundance of *Campylobacter*, with pigs from supplemented sows and offered a basal diet post-weaning having the greatest reduction. Members of the genus *Campylobacter* are commonly associated with enteritis and gastroenteritis in humans [34]. Within the phylum Spirochaetes, the abundance of the genus *Treponema* was decreased in SS pigs. *Treponema* is associated with ear necrosis and shoulder ulcers in pigs [35, 36], thus a lesser abundance is optimal. Overall, the increase in *Lactobacillus* and the reduction in *Enterobacteriaceae*, *Campylobacter* and *Treponema* suggests an overall improvement in the gut microbiome of pigs weaned from supplemented sows.

Post-weaning diarrhoea is a consequence of the reduced digestive and absorptive capacity of the small intestine which leads to the proliferation of pathogenic bacteria, mainly Enterotoxigenic *Escherichia coli* (ETEC), in the large intestine [37]. The SS pigs had the lowest faecal scores (healthiest) compared to all other groups with no evidence of the transient rise in faecal scores and diarrhoeal incidence, usually evident in the immediate post-weaning period. The period of supplementation was influential, as pigs weaned from supplemented sows had reduced diarrhoeal incidence compared to pigs weaned from basal sows and pigs offered the supplement post-weaning had reduced diarrhoea incidence compared to pigs offered the basal diet post-weaning. Pigs weaned from supplemented sows had reductions in *Enterobacteriaceae* in colonic digesta compared to pigs weaned from basal sows, perhaps contributing to the reduction in faecal scores and incidence of diarrhoea. Maternal supplementation of the CH-YBG increased the expression of *CLDN1* in the jejunum. Claudins regulate the epithelial permeability through the formation of tight junctions. Increased expression of *CLDN1* in the current study is likely to enhance the epithelial barrier during periods of challenge.

Volatile fatty acids are essential for the growth and proliferation of colonocytes [38]. They allow for greater energy utilisation in the large intestine by retrieving energy from carbohydrates not digested in the small intestine [39]. Of significance in the current study was the increased concentration of butyrate in pigs weaned from supplemented sows (SB, SS). The increase in butyrate is perhaps due to the increased abundance of *Lachnospira* in pigs weaned from supplemented sows and offered the basal diet post-weaning (SB). *Lachnospiraceae* is one of the main butyrate producing microbes in the colon [40]. Butyrate is associated with the fermentation of fibre [41] and is the favoured energy source of colonocytes accounting for approximately 70% of total energy consumption [42]. Volatile fatty acids also promote the absorption of sodium and water thus having antidiarrheal effects [42, 43] potentially contributing to the reduced diarrhoea incidence observed in pigs weaned from the supplemented sows (SB, SS).

In the current study, feed intake was reduced while gain-to-feed (feed efficiency) was improved in pigs weaned from supplemented sows (SB, SS) compared to pigs weaned from basal sows (BB, BS). The exact mechanism behind the reduced feed intake is not clear and its exploration was beyond the scope of the study. However, it is an interesting find and warrants further investigation. Maximising feed intake during the immediate post-weaning period is considered crucial in pig production as a low feed intake negatively affects nutrient uptake and utilisation [44]. A plausible explanation for the reduced feed intake is an increase in satiety due to specific milk peptides [45]. This could occur either directly or indirectly via regulatory molecules such as CCK [46, 47]. In fact, in the current study, maternal supplementation caused an up-regulation of *CCK* in the ileum. The improved feed efficiency in pigs weaned from supplemented sows could be attributed to improvements in the intestinal microbiota. McCormack et al. (48) identified a profile of intestinal microbiota associated with improved gut function, digestion and improved feed efficiency, with efficient pigs having an increased abundance of *Lactobacillus* which is evident in the current study. Similarly, Vigors et al. (49) reported a correlation between high feed efficiency and increased abundance of *Lactobacillus* in the caecum of pigs. The reduced feed intake corresponds with the downregulation of glucose transporters *SLC2A2/GLUT2, SLC2A5/GLUT5* and *SLC5A1/SGLT1*, peptide transporter *SLC15A1/PEPT1* and long-chain fatty acid transporter *FABP2* in the ileum in pigs weaned from supplemented sows. It is plausible that the reduced feed intake observed in the current study led to the downregulation of nutrient transporter gene expression.

## 5. Conclusion

In conclusion, a combination of both maternal and direct supplementation with CH-YBG is the desirable feeding strategy to ameliorate the post-weaning challenge. Maternal supplementation increased feed efficiency and butyrate concentrations, while also increasing the abundance of *Lactobacillus* and decreasing the abundance of *Enterobacteriaceae* and *Campylobacteraceae*. The combination of maternal and direct supplementation led to pigs having the lowest/healthiest faecal scores compared to all other groups with no evidence of the transient rise in faecal scores and diarrhoeal incidence that is commonly evident in the immediate post-weaning period.

## Ethics Statement

The authors confirm that the ethical policies of the journal, as noted on the journal’s author guidelines page, have been adhered to and the appropriate ethical review committee approval has been received. The authors confirm that they have followed the EU standards for the protection of animals used for scientific purposes.

## Acknowledgements

This research was a part of Food for Health Ireland (www.fhi.ie) project funded by Enterprise Ireland (Grant Number CC200800001)

## Author Contribution

The author’s contributions were as follows: T.S. and J.V.O.D. designed the experiment and supervised data collection; A.M and S.M performed the experiment and collected the samples; E.C carried out the laboratory analyses and wrote the manuscript; A.D., and M.T.R. contributed to laboratory analyses. T.S., J.V.O.D., and S.V. performed the statistical analyses and corrected the manuscript; All authors approved the final version of the manuscript.

